# Design of potent membrane fusion inhibitors against SARS-CoV-2, an emerging coronavirus with high fusogenic activity

**DOI:** 10.1101/2020.03.26.009233

**Authors:** Yuanmei Zhu, Danwei Yu, Hongxia Yan, Huihui Chong, Yuxian He

**Author notes:** These authors contributed equally to this work (Yuanmei Zhu, Danwei Yu). Correspondence should be addressed to Y.H., Chinese Academy of Medical Sciences, Beijing 100730, China.

## Abstract

The coronavirus disease COVID-19, caused by emerging SARS-CoV-2, has posed serious threats to global public health, economic and social stabilities, calling for the prompt development of therapeutics and prophylactics. In this study, we firstly verified that SARS-CoV-2 uses human ACE2 as a cell receptor and its spike (S) protein mediates high membrane fusion activity. Comparing to that of SARS-CoV, the heptad repeat 1 (HR1) sequence in the S2 fusion protein of SARS-CoV-2 possesses markedly increased α-helicity and thermostability, as well as a higher binding affinity with its corresponding heptad repeat 2 (HR1) site. Then, we designed a HR2 sequence-based lipopeptide fusion inhibitor, termed IPB02, which showed highly poent activities in inibibiting the SARS-CoV-2 S protein-mediated cell-cell fusion and pseudovirus infection. IPB02 also inhibited the SARS-CoV pseudovirus efficiently. Moreover, the strcuture and activity relationship (SAR) of IPB02 were characterzized with a panel of truncated lipopeptides, revealing the amino acid motifs critical for its binding and antiviral capacities. Therefore, the presented results have provided important information for understanding the entry pathway of SARS-CoV-2 and the design of antivirals that target the membrane fusion step.

## Introduction

In late December of 2019, a new infectious respiratory disease emerged in Wuhan, China. The pathogen was soon identified as a novel coronavirus (CoV) (1-3), which was initially termed 2019-nCoV by the World Health Organization (WHO) and the disease was named COVID-19 (2019 Coronavirus Disease). Because 2019-nCoV shares a high sequence identity to the previously emerged severe acute respiratory syndrome CoV (SARS-CoV) and the same cell receptor angiotensin-converting enzyme 2 (ACE2) for infection, it was renamed SARS-CoV-2 by the Coronaviridae Study Group (CSG) of the International Committee on Taxonomy of Viruses (ICTV). As of 26 March 2020, a total of 416,686 confirmed COVID-19 cases, including 18,589 deaths, have been reported from 197 countries or regions (www.who.int/emergencies/diseases/novel-coronavirus-2019). The pandemic has posed serious threats to global public health, economic and social stabilities, calling for the urgent development of vaccines and antiviral drugs.

CoVs, a large group of enveloped viruses with a single positive-stranded RNA genome, are genetically classified into four genera: α-, β-, γ-, and δ-CoVs (4, 5). The previously known six CoVs that cause human disease include two α-CoVs (NL63; 229E) and four β-CoVs (OC43; HKU1; SARS-CoV; MERS-CoV). SARS-CoV-2 belongs to the β-CoV genus and represents the seventh human CoV. Like other CoVs, SARS-CoV-2 use a glycosylated, homotrimeric class I fusion spike (S) protein to gain entry into host cells (6-8). The S protein comprises of S1 and S2 subunits and exists in a metastable prefusion conformation. The S1 subunit, which contains a receptor-binding domain (RBD) capable of functional folding independently, is responsible for virus binding to the cell surface receptor. A recent study suggested that ACE2-binding affinity of the RBD of SARS-CoV-2 is up to 20-fold higher than that of SARS-CoV, which may contribute to the significantly increased infectivity and transmissibility (6). The receptor-binding deem to trigger large conformational changes in the S complex, which destabilize the prefusion trimer resulting in shedding of the S1 subunit and activate the fusogenic activity of the S2 subunit (9-11). As illustrated in Fig. 1, the sequence structure of S2 contains an N-terminal fusion peptide (FP), heptad repeat 1 (HR1), heptad repeat 2 (HR2), transmembrane region (TM), and cytoplasmic tail (CT). During the fusion process, the FP is exposed and inserts into the target cell membrane, leading S2 in a prehairpin intermediate that bridges the viral and cell membranes; then, three HR1 segments self-assemble a trimeric coiled-coil and three HR2 segments fold into the grooves on the surface of the HR1 inner core, thereby resulting a six-helical bundle (6-HB) structure that drives the two membranes in close apposition for fusion.

**Figure 1.**
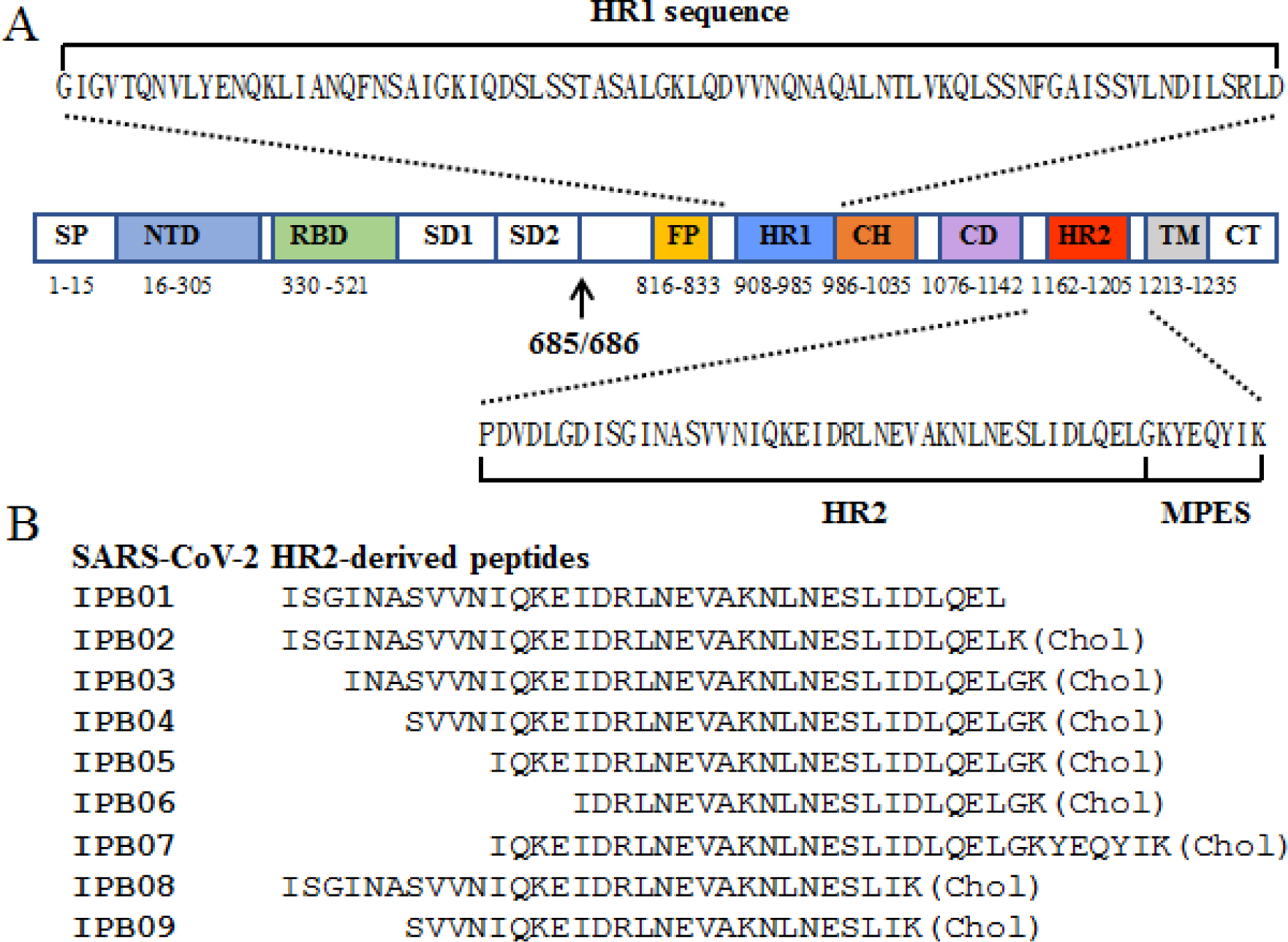
Schematic diagram of SARS-CoV-2 S protein and its peptide derivatives. (**A**) Functional domains of the S protein. SP, signal peptide; NTD, N-terminal domain; RBD, receptor-binding domain; SD, subdomain; FP, fusion peptide; HR1, heptad repeat 1; CH, central helix; CD, connector domain; HR2, heptad repeat 2; TM, transmembrane domain; CT, cytoplasmic tail. The S1/S2 cleavage site (685/686) is marked. The HR1 and HR2 sequences and membrane proximal external sequence (MPES) are listed. (**B**) HR2-derived fusion inhibitor peptides. chol, cholesterol.

Peptides derived from the HR1 and HR2 sequences of the class I viral fusion proteins have been demonstrated to possess antiviral activity through binding to the prehairpin intermediate thus blocking the formation of viral 6-HB core (12). Such are indeed the cases for emerging CoVs, including SARS-CoV and MERS-CoV (10, 13-15). In response to the outbreak of SARS-CoV, a group of HR2-based peptides that could effectively inhibit viral infection were developed (10, 15-18). Recently, a pan-CoV fusion inhibitor, designated EK1, was created, which showed inhibitory activities against diverse HCoVs, including SARS-CoV, MERS-CoV, HCoV-229E, HCoV-NL63, and HCoV-OC43 (19). However, the previously reported fusion inhibitor peptides often display low antiviral activities, with a 50% inhibitory concentration (IC_50_) at macromolar (μM) range. In the past decade, we have dedicated our efforts to develop viral fusion inhibitors with improved pharmaceutical profiles, generating a group of lipopeptides with extremely potent antiviral activity (20-25). To fighting the COVID-19 pandemic, here we have applied our expertise to develop fusion inhibitors against SARS-CoV-2 infection. We found that different from that of SARS-CoV, the S protein of SARS-CoV-2 has a high cell fusion activity; then, we designed and characterized several lipopeptide-based fusion inhibitors with highly potent activities in inhibiting both SARS-CoV-2 and SARS-CoV.

## Results

### SARS-CoV-2 uses ACE2 as a cell receptor and its S protein displays high fusion activity

In the earlier time point, we would like to experimentally verify whether SARS-CoV-2 uses human ACE2 as a receptor for cell entry, thus we generated its S protein pseudotyped lentiviral particles. The SARS-CoV and vesicular stomatitis virus (VSV-G) pseudoviruses were also prepared for comparison. As shown in Fig. 2A, all of three pseudoviruses efficiently infected 293T cells that stably overexpress ACE2 (293T/ACE2); however, the infectivity SARS-CoV-2 and SARS-CoV dramatically decreased in 239T cells which express a low level of endogenous ACE2. As a virus control, VSV-G pesudovirus entered 239T cells even more efficiently relative its infectivity in 293T/ACE2 cells.

**Figure 2.**
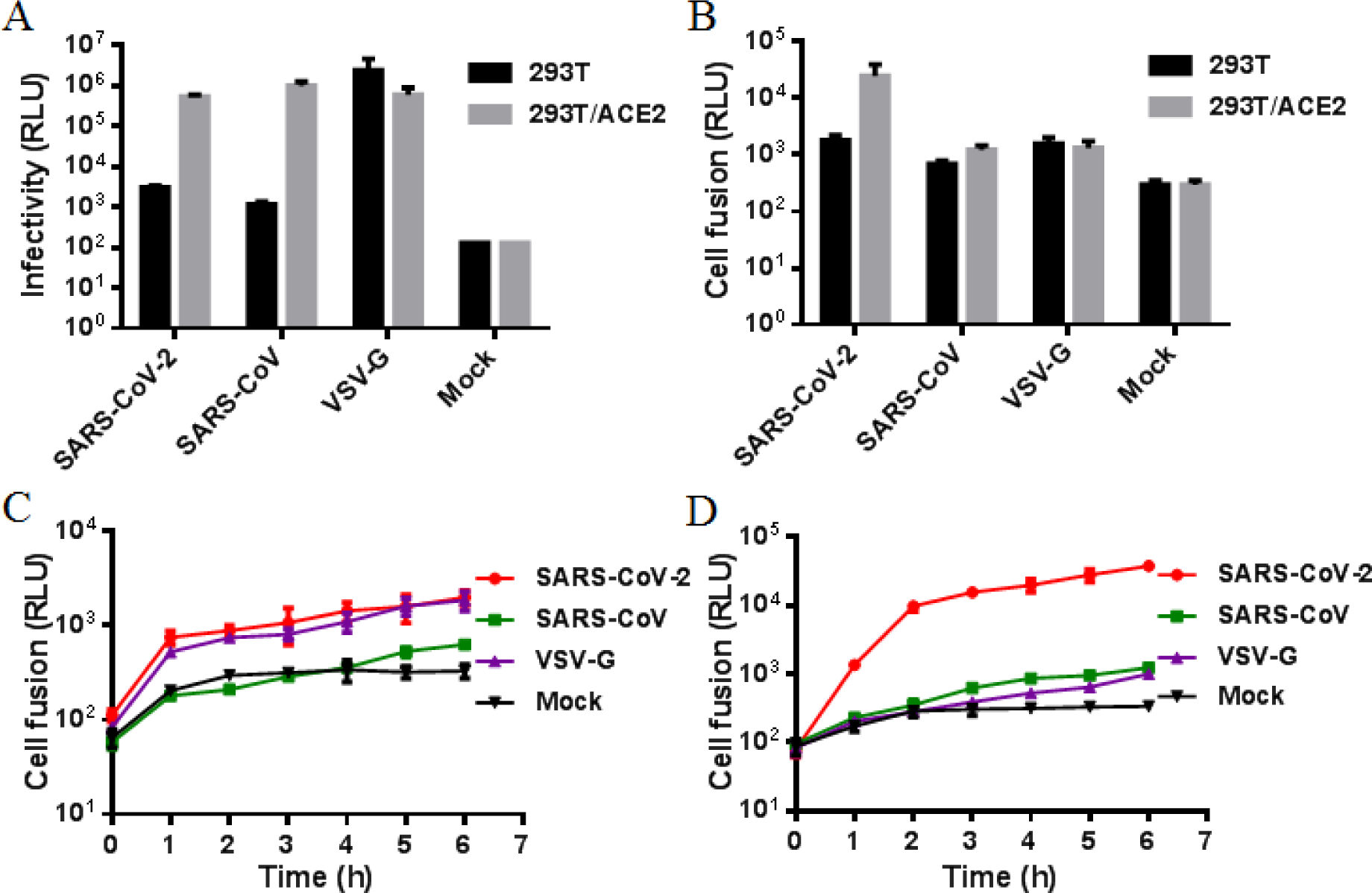
Functional characterization of the SARS-CoV-2 and SARS-CoV S proteins. (**A**) Infectivity of the SARS-CoV-2 and SARS-CoV pseudoviruses in 293T cells or 293T/ACE2 cells was determined by a single-cycle infection assay. (**B**) Fusogenic activity of the SARS-CoV-2 and SARS-CoV S proteins with 293T cells or 293T/ACE2 cells as a target was determined by a DSP-based cell fusion assay. The fusion activity of S proteins in 293T cells (**C**) and 293T/ACE2 cells (**D**) were determined at different time points. The experiments were repeated three times, and data are expressed as means ± standard deviations.

We further compared the fusion activity of viral S protein in 293T and 293T/ACE2 cells by applying a DSP-based cell-cell fusion assay. As shown in Fig. 2B, both the S proteins of SARS-CoV-2 and SARS-CoV displayed a weak fusion activity in 293T cells, but they showed significantly increased capacities to mediate cell fusion with 293T/ACE2 cells. These results demonstrated that overexpression of ACE2 can promote the cell entry of both the SARS-CoV-2 and SARS-CoV pseudoviruses as well as the S protein-mediated cell-cell fusion activity, verifying the functionality of ACE2 for SARS-CoV-2.

In both the 239T and 293T/ACE2 target cells, we observed that the S protein of SARS-CoV-2 had a significantly increased fusion activity than the S protein of SARS-CoV. Therefore, we further compared the fusion activities of viral S proteins at different time points. As shown in Fig. 2C and 2D, the SARS-CoV S protein exhibited had no appreciable fusion activity until the effector cells and target cells were cocultured for five or six hours; in sharp contrast, the SARS-CoV-2 S protein mediated a rapid and robust cell fusion reaction, as indicated by its fusion kinetic curves especially in 293T/ACE2 cells.

### Compared to SARS-CoV, SARS-CoV-2 might possess an enhanced HR1-HR2 interaction

Similar to many class I fusion proteins, the interaction between the HR1 and HR2 domains of the CoV fusion protein S2 critically determines viral membrane fusion activity. Comparing to SARS-CoV, SARS-CoV-2 has a HR1 sequence with nine amino acid substitutions, and of them eight are located within the HR1 core site; whereas, two viruses share a fully identical HR2 sequence (Fig. 3A). In order to explore the mechanism underlying the highly active fusion activity of the SARS-CoV-2 S protein, we synthesized two peptides corresponding to the HR1 sequence and their secondary structures were determined by circular dichroism (CD) spectroscopy. As shown in Fig. 3B, the HR1 peptide derived from SARS-CoV-2, designated SARS2NP, showed a typical α-helical conformation with the helix contents of 66%, whereas the HR1 peptide from SARS-CoV, designated SARS1NP, had α-helical contents of 41%. The thermal stability of the two peptides was further measured. As shown in Fig. 3C, SARS2NP and SARS1NP exhibited their melting temperature (*T*_m_) values of 48 and 40°C, respectively. Furthermore, we synthesized a peptide containing the HR2 sequence, termed IPB01, and its interactions with the two HR1 peptides were analyzed by CD spectroscopy. As shown in Fig. 3D and E, both the SARS2NP and SARS1NP interacted with IPB01 to form complexes with typical α-helical structures, having the *T*_m_ values of 75 and 68 °C, respectively. In comparison, the complex formed by SARS2NP and IPB01 was much more stable than the complex between the SARS1NP and IPB01 peptides. Taken together, these results suggested that SARS-CoV-2 might evolve an increased interaction between the HR1 and HR2 domains in the S2 fusion protein thus critically determining its high fusogenic activity.

**Figure 3.**
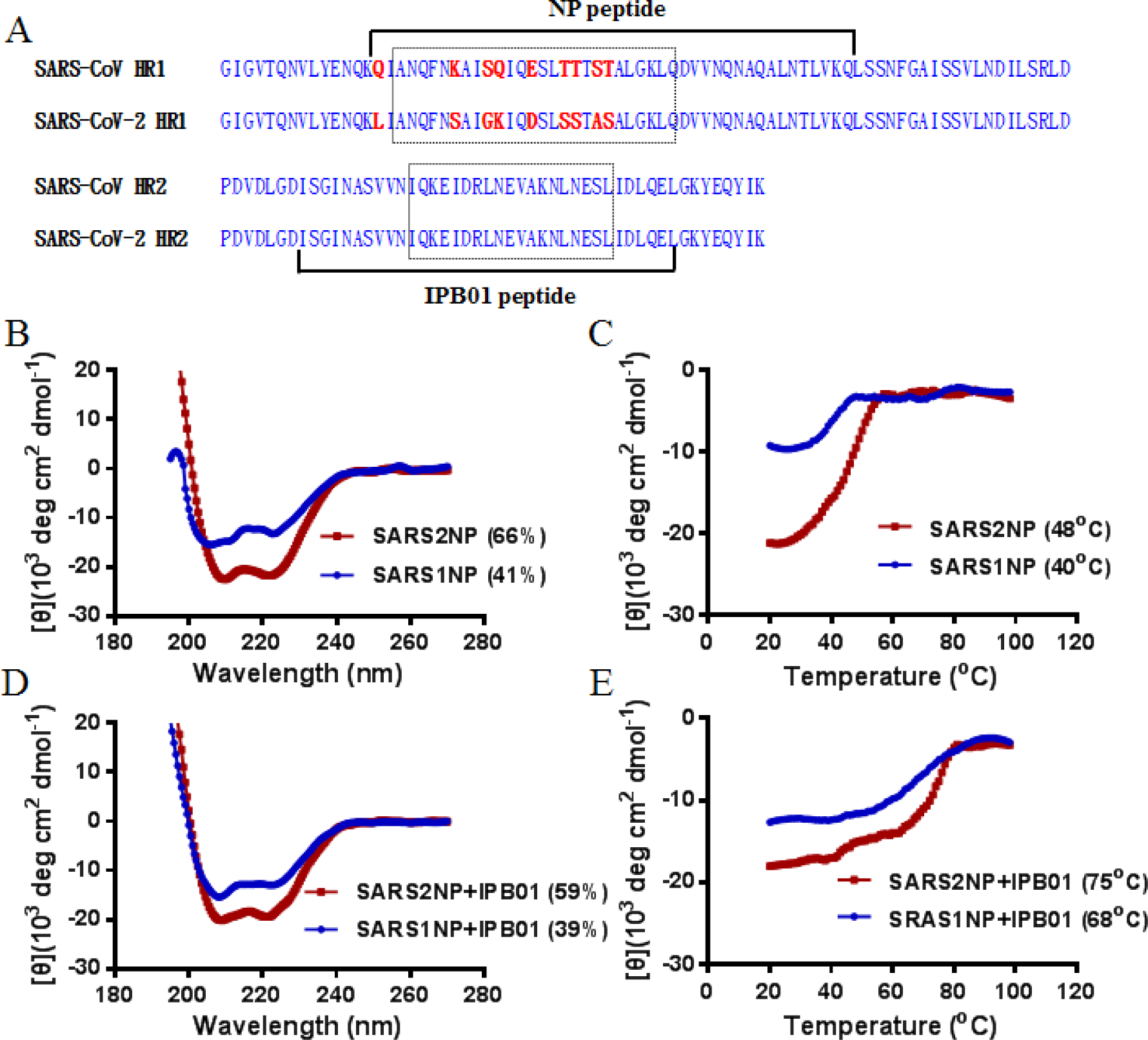
The interactions between the HR1 and HR2 peptides derived from the S2 proteins of SARS-CoV-2 and SARS-CoV. (**A**) Sequence comparison of the HR1 and HR2 domains in SARS-CoV-2 and SARS-CoV. The α-helicity and thermostability of the HR1 peptides alone (**B** and **C**) or in complexes with a HR2 peptide (**D** and **E**) were determined by CD spectroscopy, in which the peptides or peptide mixture were dissolved in PBS with a final concentration of each peptide at 10 μM. The experiments were performed two times, and representative data are shown.

### Cholesterylated peptide exhibits greatly increased α-helical stability and target-binding affinity

Emerging studies demonstrate that lipid conjugation is a viable strategy to design peptide-based viral fusion inhibitors with enhanced antiviral activity and *in vivo* stability. The resulting lipopeptides are considered to interact preferentially with the viral and cell membranes, thus raising the local concentration of the inhibitors at the site where viral fusion occurs (20-25). According to our previous experiences, here we modified the HR2 peptide IBP01 by adding a cholesterol group to its C-terminal, resulting in a lipopeptide termed IPB02, as illustrated in Fig. 1B. We first applied CD spectroscopy to determine the structural properties of the inhibitors in the absence or presence of a target mimic HR1 peptide. As shown in Fig. 4A and 4B, the unconjugated IPB01 alone was largely in a random structure and its *T*_m_ value could not be defined. By contrast, the lipopeptide IPB02 displayed markedly increased helix contents with a *T*_m_ of 65 °C. Next, we assessed the helical binding stability of the inhibitors with the two target mimic peptides, SARS2NP and SARS1NP. As shown in Fig. 4C to 4F, the lipopeptide-based complexes had sharply increased thermostabilities compared to the complexes formed by the template peptides. Specifically, the IPB02 and SARS2NP complex showed a *T*_m_ of 89°C, which was 14°C higher than the IPB01 and SARS2NP complex (75°C); the IPB02 and SARS1NP complex also had a *T*_m_ of 89°C, indicating a 21°C increase relative to the IPB0-based complex (68°C). Here the CD results demonstrated that the cholesterol conjugated peptide IPB02 possesses significantly increased α-helical thermostability and target-binding affinity.

**Figure 4.**
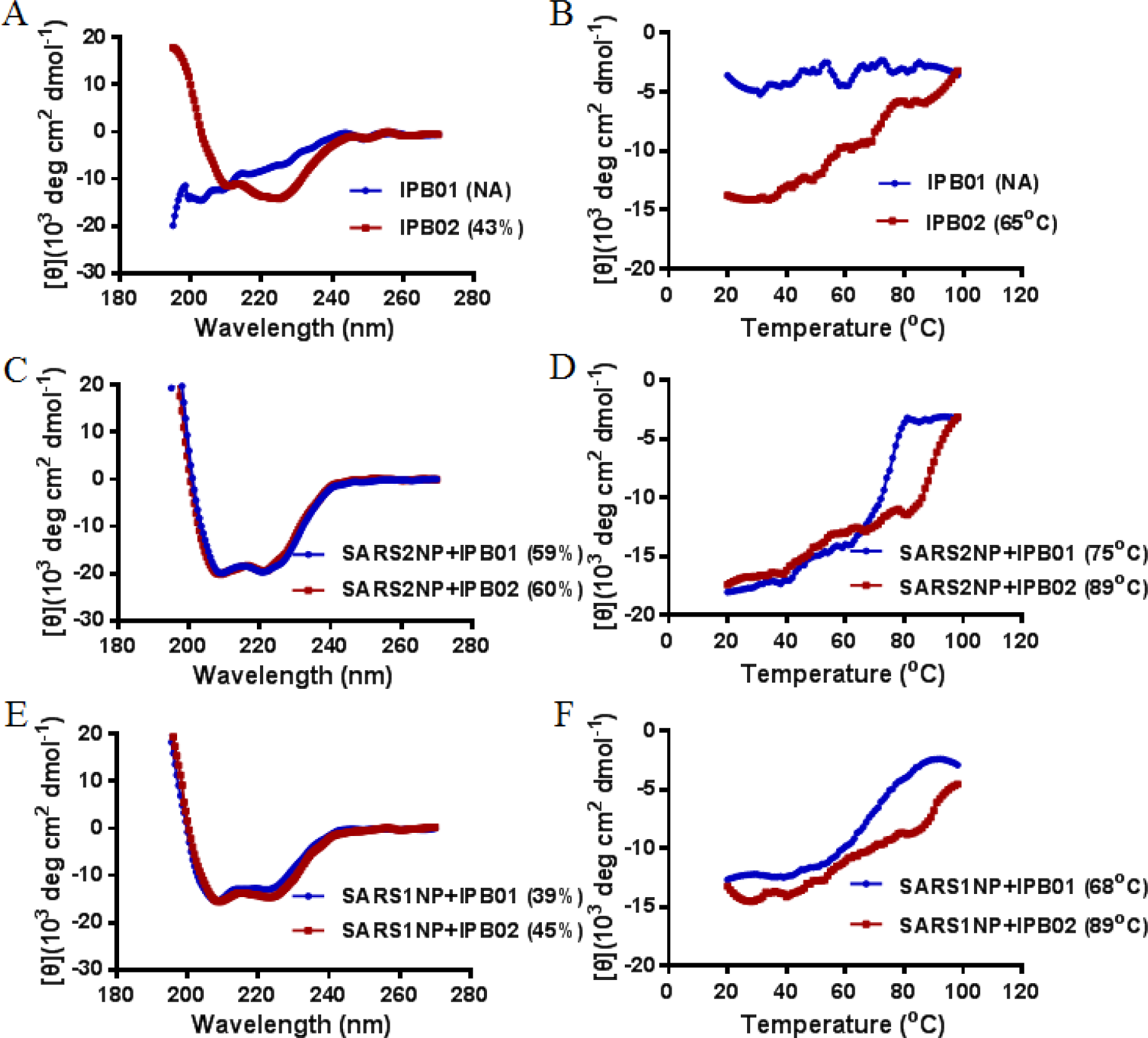
Secondary structure and binding stability of fusion inhibitor peptides determined by CD spectroscopy. The α-helicity and thermostability of peptide inhibitors alone (**A** and **B**) or in complexes with the SARS-CoV-2 HR1 peptide (**C** and **D**) or SARS-CoV HR1 peptide (**E** and **F**) were detected with a final concentration of each peptide at 10 μM. The experiments were performed two times, and representative data are shown.

We also visualized the formed complexes by a native-polyacrylamide gel electrophoresis (PAGE) method. As shown in Fig. 5, the positively charged SARS2NP and SARS1NP might migrate up and off the gel thus no bands appeared, whereas IPB01 and IPB02 showed specific bands because they carried net negative charges. When a HR1 peptide and an inhibitor were mixed, new bands corresponding to the binding complexes emerged at the upper positions of the gel, which verified the between interactions.

**Figure 5.**
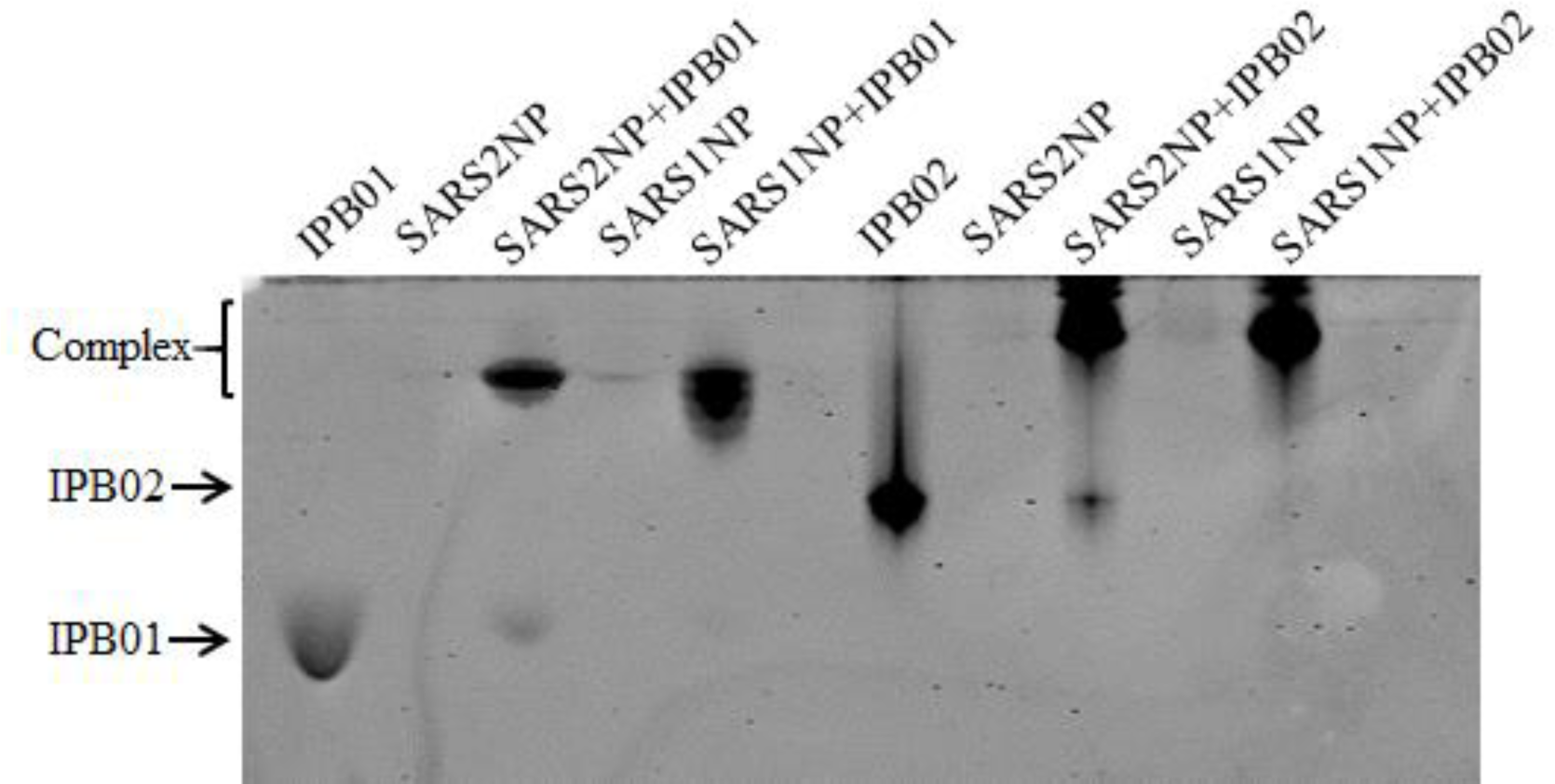
Visualization of the interactions between HR1 peptides and inhibitors by native PAGE analysis. Each of the peptides was used at a final concentration of 40μM. The positively charged peptides SARS2NP and SARS1NP migrated up and off the gel, thus no bands appeared. IPB01 or IPB02 alone and their complexes with SARS2NP or SARS1NP displayed specific bands because of their net negative charges. The experiments were repeated two times, and representative data are shown.

### IPB02 is a highly potent fusion inhibitor of SARS-CoV-2 and SARS-CoV

We next sought to determine the antiviral functions of the IPB01 and IPB02 peptides. Firstly, their inhibitory activities on S protein-mediated cell-cell fusion were examined by the DSP-based cell fusion assay as described above. As shown in Fig. 6A and Table 1, both of IPB01 and IPB02 potently inhibited the cell fusion mediated by the S protein of SARS-CoV-2, with mean IC_50_ values of 0.022 and 0.025 μM, respectively. Then, we conducted the single-cycle infection assay to measure the inhibitory activities of the peptides on pseudoviruses. Surprisingly, the unconjugated peptide IPB01 showed very weak or marginal activities in inhibiting the SARS-CoV-2 (Fig. 6B) and SARS-CoV (Fig. 6C) pseudoviruses; however, the lipopeptide IPB02 inhibited the two viruses with IC_50_ at 0.08 and 0.251μM, respectively (Table 1). As expected, IPB01 and IPB02 had no inhibitory activity against a control virus (VSV-G), indicating their antiviral specificities. Therefore, we conclude that IPB02 is a highly potent fusion inhibitor of SARS-CoV-2 and SARS-CoV.

**Table 1.** Structural and functional characterization of lipopeptide fusion inhibitors against SARS-CoV-2 and SARS-CoV

| Inhibitor | Inhibitory activity (IC<sub>50</sub>, μM) | | | | Helix content (%) | | $T_m$ (°C) | |
| --- | --- | --- | --- | --- | --- | --- | --- | --- |
|  | SARS-CoV-2 fusion | SARS-CoV-2 PV | SARS-CoV PV | VSV-G PV | SARS2NP | SARS1NP | SARS2NP | SARS1NP |
| IPB01 | 0.022 ± 0.005 | 33.74 ± 11.827 | >50 | >50 | 59 | 39 | 75 | 68 |
| IPB02 | 0.025 ± 0.002 | 0.08 ± 0.017 | 0.251 ± 0.118 | >50 | 60 | 45 | 89 | 89 |
| IPB03 | 0.015 ± 0.002 | 0.947 ± 0.179 | 1.315 ± 0.463 | >50 | 50 | 34 | 47 | 46 |
| IPB04 | 0.033 ± 0.013 | 0.218 ± 0.063 | 1.053 ± 0.444 | >50 | 60 | 49 | 77 | 77 |
| IPB05 | >5 | >25 | >25 | >50 | ND | ND | ND | ND |
| IPB06 | >5 | >25 | >25 | >50 | ND | ND | ND | ND |
| IPB07 | 0.017 ± 0.001 | 0.993 ± 0.08 | 1.037 ± 0.836 | >50 | 55 | 33 | 47 | 45 |
| IPB08 | 4.66 ± 1.565 | 1.738 ± 0.898 | 1.13 ± 0.472 | >50 | ND | ND | ND | ND |
| IPB09 | >5 | >25 | >25 | >50 | ND | ND | ND | ND |

\*The antiviral assays were repeated three times, and data are expressed as means ± standard deviations. The CD experiment was repeated two times and representative data are shown. ND means “not done” owing to the solubility problem of the peptides in PBS.

**Figure 6.**
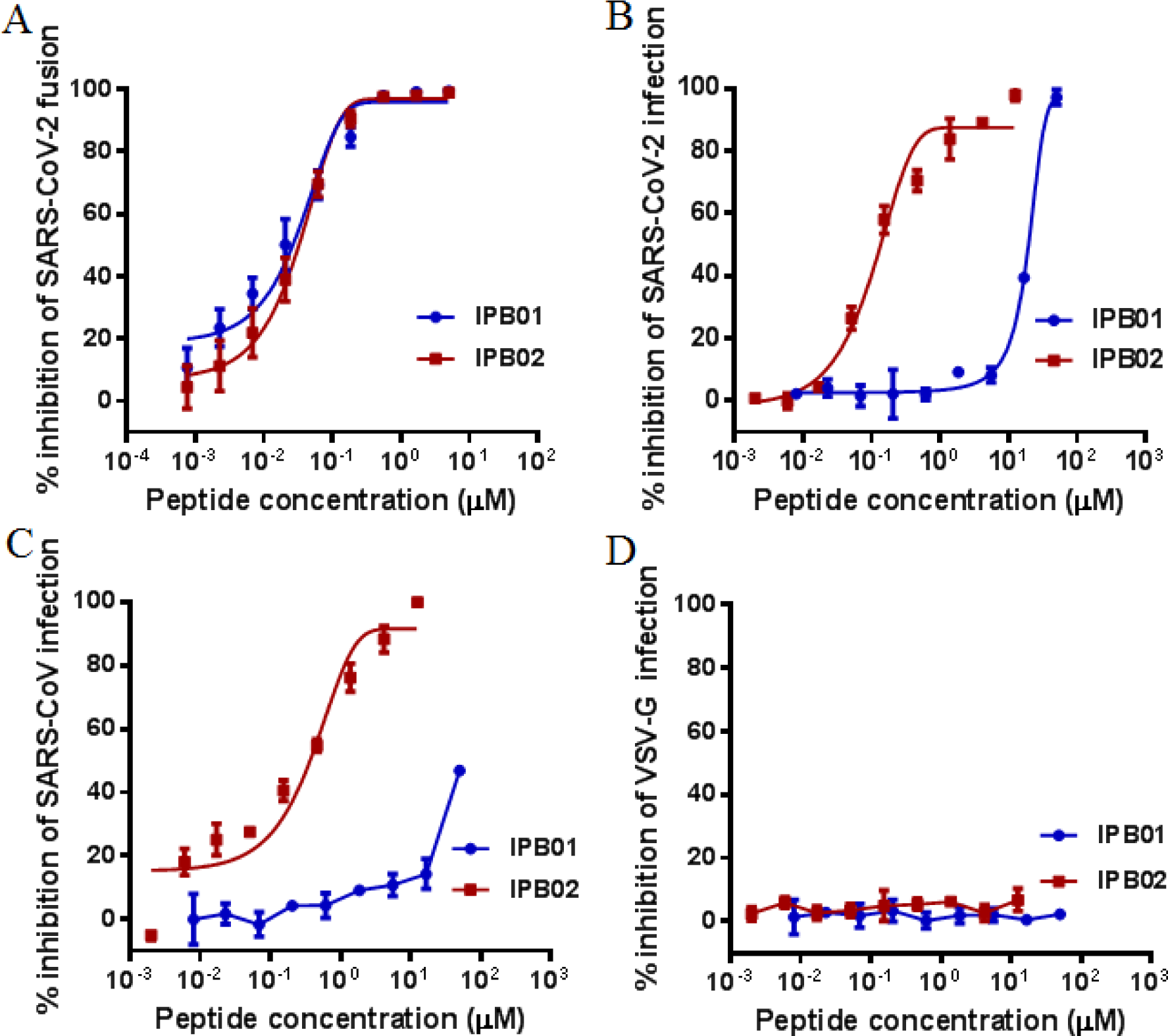
Inhibitory activity of IPB01 and IPB02 against SARS-CoV-2 and SARS-CoV. (**A**) Inhibition of inhibitors on the SARS-CoV-2 S protein-mediated cell-cell fusion determined by a DSP-based cell fusion assay. The activity of IPB01 and IPB02 in inhibiting SARS-CoV-2 (**B**) and SARS-CoV (**C**) and a control pseudovirus VSV-G (**D**) was determined by a single-cycle infection assay. The experiments were repeated three times, and data are expressed as means ± standard deviations.

### Structural and functional characterization of lipopeptide inhibitors

In light of the high binding and inhibitory activities with IPB02, we next focused on characterizing its structure-activity relationship (SAR). To this end, a panel of new lipopeptides was generated by sequence truncation or extension, and their antiviral capacities were examined. As shown in Table 1, IPB03 and IPB04, which had an N-terminal amino acid truncation, still maintained a very high potency in inhibiting the cell fusion activity of the SARS-CoV-2 S protein, but they exhibited an obviously reduced activity to block the cell entry activity of both the SARS-CoV-2 and SARS-CoV pseudoviruses. A further N-terminal truncation, as indicated by IPB05 and IPB06, would result in the inhibitors inactive at a high concentration. By adding six amino acids of the membrane proximal external sequence (MPES) to the C-terminal of IPB05, the resulting peptide IPB07 regained the antiviral activity, demonstrating the importance of MPES in the design of such CoV fusion inhibitors. Differently, IPB08 was a C-terminally truncated inhibitor with IPB02 as a template, but its antiviral function was markedly impaired, underscoring the roles of C-terminal residues in IPB02. On the basis of the results above, it was expected that IPB09 with two terminal truncations was antivirally inactive. Indeed, the CD data suggested that both the N- and C-terminal sequences contributed critically to the binding of the inhibitors (Table 1). By comparing IPB03 and IPB04, it revealed that three amino acids (Ile-Asn-Ala) in the N-terminal of IPB03 reversely impaired the inhibitor binding.

## Discussion

In 2002, SARS-CoV suddenly emerged in Guangzhou, China, and its subsequent global spread was associated with 8096 cases and 774 deaths. To fight against SARS-CoV, we took immediate actions with multiple research projects and achieved significant findings. First, we identified several viral antigens suitable for the development of diagnostic tools (26-28); second, we proposed for the first time that the S protein receptor-binding domain (RBD) can serve as an ideal subunit vaccine for emerging CoVs (29-40); third, we also reported the first peptide-based SARS-CoV fusion inhibitor with potential therapeutic and preventive efficacies (10).

To fight against the current pandemic of COVID-19 caused by SARS-CoV-2, we sprang into action to develop effective therapeutics and prophylactics. In this study, we focused on developing viral fusion inhibitor peptides with a potent and broad antiviral activity. Firstly, our experiments verified that like SARS-CoV, SARS-CoV-2 also uses human angiotensin-converting enzyme 2 (ACE2) as a receptor for cell entry and infection; however, the S protein of SARS-CoV-2 has much higher activity to mediate cell-cell fusion. By analyzing the secondary structure and thermostability with CD spectroscopy, we found that the HR1 peptide derived from the S2 fusion protein of SARS-CoV-2 displays much higher α-helicity and thermostability than the HR1 peptide from the S2 fusion protein of SARS-CoV. Consistently, both the α-helical contents and melting temperature of the SARS-CoV-2 HR1 peptide complexed with a HR2-derived peptide are higher than that of the SARS-CoV HR1 peptide-based complex, suggesting a more strong interaction between the HR1 and HR2 sites for SARS-CoV-2 over that of SARS-CoV. According to our experiences in designing of lipopeptide fusion inhibitor against HIV, we modified the HR2 sequence-derived peptide IPB01 with a cholesterol group, resulting in the lipopeptide IPB02 with highly potent activities in inhibiting SARS-CoV-2 and SARS-CoV pseudoviruses as well as the S protein-mediated cell-cell fusion activity. Moreover, the structure-activity relationship (SAR) of the HR2 sequence-based fusion inhibitors were characterized by applying a panel of truncated lipopeptides, which certified the roles of both the N-and C-terminal amino acid sequences in the design of a potent inhibitor against emerging CoVs. Combined, these data provide important information for understanding the fusion mechanism of emerging CoVs and for the development of antivirals that target the membrane fusion step.

Cell entry of CoVs depends on binding of the viral S proteins to cellular receptors and on S protein priming by host cell proteases (41-44). Previous studies demonstrated that SARS-CoV enters into targeting cells mainly via an endosome membrane fusion pathway where its S protein is cleaved by endosomal cysteine proteases cathepsin B and L (CatB/L) and activated (45). However, SARS-CoV also employs the cellular serine protease TMPRSS2 for S protein priming, and especially, TMPRSS2 but not CatB/L is essential for viral entry into primary target cells and for viral spread in the infected host (43, 46-48). It was also found that introducing a furin-recognition site between the S1 and S2 subunits could significantly increase the ability of SARS-CoV S protein to mediate cellular membrane surface infection (49). Differently, SARS-CoV-2 induces typical syncytium formation in infected cells, suggesting that it mainly utilizes a plasma membrane fusion pathway for cell entry. Sequence analyses revealed that SARS-CoV-2 harbors the S1/S2 cleavage site in its S protein, although its roles in S protein-mediated membrane fusion and viral life-cycle need to be characterized. One can speculate that furin-mediated precleavage at the S1/S2 site in infected cells might promote subsequent TMPRSS2-dependent cell entry, the case for MERS-CoV (50, 51). A recent study found that SARS-CoV-2 employs TMPRSS2 for S protein priming and a TMPRSS2 inhibitor approved for clinical use can block entry (52). For most viruses, the plasma membrane fusion pathway is more efficient than the endosome membrane fusion pathway because the latter is prone to activating the host antiviral immunity (53, 54). In this study, we have not only verified ACE2 as a cell receptor but also demonstrated that the SARS-CoV-2 S protein evolves a significantly increased fusogenic activity relative to the S protein of SARS-CoV. Although our studies have also shown that the HR1 mutations in the S2 protein can greatly enhance the HR1-HR2 interaction thus might be a crucial factor to determine the fusion activity of the SARS-CoV-2 S protein, other players in the fusion pathways may contribute in coordination with that. We also observed that the HR2-derived inhibitors were more effective in inhibiting the S protein-mediated cell fusion than their inhibitions on pseudoviruses, implying that SARS-CoV-2 might also adopt the endosome entry pathway. Indeed, it was recently found that cathepsin L is required for the cell entry of SARS-CoV-2 and teicoplanin, a glycopeptide antibiotic, can specifically inhibit the entry (55, 56).

Drug repurposing represents as a viable drug discovery strategy from existing drugs with knowledge on safety profile, side effects, posology and drug interactions, which could shorten the time and reduce the cost compared to *de novo* drug discovery. In the emergency of the COVID-19 pandemic, a group of nonspecific antiviral drugs, including interferon (IFN), lopinavir/ritonavir, chloroquine, remdesivir (GS-5734), and favipiravir (T-705) were quickly screened with anti-SARS-CoV-2 activity and they have been used to treat infected patients (57, 58). Very recently, a clinical study suggested that combination of hydroxychloroquine and azithromycin would provide synergistic effects in treated COVID-19 patients (59). Nonetheless, development of specific drugs for emerging coronaviruses, including SARS-CoV-2 and SARS-CoV, is highly required and has perspective for the long run. We believe that the newly developed lipopeptide IPB02 represents an ideal candidate for future optimization development.

## Materials and methods

### Peptide synthesis

Peptides were synthesized on rink amide 4-methylbenzhydrylamine (MBHA) resin using a standard solid-phase 9-flurorenylmethoxycarbonyl (FMOC) protocol as described previously (20). Lipopeptides were produced by conjugating cholesterol succinate monoester to the C-terminal lysine residue. All peptides were N-terminally acetylated and C-terminally amidated, and they were purified by reverse-phase high-performance liquid chromatography (HPLC) to more than 95% homogeneity and characterized with mass spectrometry.

### Single-cycle infection assay

Infectivity of SARS-CoV-2, SARS-CoV, and vesicular stomatitis virus (VSV) on 293T cells or 293T cells stably expressing human ACE2 (293T/ACE2) was determined by a single-cycle infection assay as described previously (60). To produce pseudoviruses, HEK293T cells were cotransfected with a backbone plasmid (pNL4-3.luc.RE) that encodes an Env-defective, luciferase reporter-expressing HIV-1 genome and a plasmid expressing the S protein of SARS-CoV-2 or SARS-CoV or the G protein of VSV. Cell culture supernatants containing the released virions were harvested 48 h post-transfection, filtrated and stored at -80°C. To measure the inhibitory activity of peptide inhibitors, pseudoviruses were mixed with an equal volume of a serially 3-fold diluted peptide and incubated at 37 °C for 30 min. The mixture was then added to 293T/ACE2 cells at a density of 10^4^ cells/100 μl per plate well. After cultured at 37 °C for 48 h, the cells were harvested and lysed in reporter lysis buffer, and luciferase activity was measured using luciferase assay reagents and a luminescence counter (Promega, Madison, WI, USA). The 50% inhibitory concentration (IC_50_) was calculated as the final cell culture concentration of an inhibitor that caused a 50% reduction in relative luminescence units (RLU) compared to the level of the virus control subtracted from that of the cell control.

### Cell-cell fusion assay

A dual split-protein (DSP)-based fusion cell-cell assay was used to detect the SARS-CoV-2 or SARS-CoV S protein-mediated cell-cell fusion activity and the inhibitory activity of peptides as described previously (60). Briefly, a total of 1.5×10^4^ 293T cells (effector cells) were seeded in a 96-well plate and 1.5×10^5^/mL of 293T or 293T/ACE2 cells (target cells) were seeded in a 10-cm culture dish, and then incubated at 37°C. On the next day, effector cells were cotransfected with a S protein-expressing and a DSP_1-7_ plasmid, target cells were transfected with a DSP_8-11_ plasmid, and then the cells were incubated at 37°C. After 24 h, the effector cells were added with a serially 3-fold diluted peptide and incubated for 1 h; the target cells were resuspended at 3×10^5^/mL in prewarmed culture medium that contains EnduRen live cell substrate (Promega) at a final concentration of 17 ng/ml and incubated for 30 min. Then, 3×10^4^ of target cells were then transferred to the effector cells and the mixed cells were spun down to maximize cell-cell contact. After incubation for 2 h, luciferase activity was measured and IC_50_ values were calculated as described above.

### Circular dichroism (CD) spectroscopy

The secondary structure and thermal stability of peptides or peptide complexes were determined by CD spectroscopy as described previously (60). Briefly, a peptide was dissolved in phosphate-buffered saline (PBS; pH 7.2) with a final concentration of 10 μM and incubated at 37 °C for 30 min. CD spectra were acquired on a Jasco spectropolarimeter (model J-815) using a 1 nm bandwidth with a 1 nm step resolution from 195 to 270 nm at room temperature. Spectra were corrected by subtracting a solvent blank, and α-helical content was calculated from the CD signal by dividing the mean residue ellipticity [θ] at 222 nm, with a value of -33,000 deg cm^2^ dmol^-1^ corresponding to 100% helix. Thermal denaturation was conducted by monitoring the ellipticity change at 222 nm from 20 to 98°C at a rate of 2°C/min, and melting temperature (*T*_*m*_) was defined as the midpoint of the thermal unfolding transition.

### Native-polyacrylamide gel electrophoresis (N-PAGE)

N-PAGE was performed to determine the interaction between a SARS-CoV2 or SARS-CoV S protein HR1-derived peptide and a HR2-derived peptide as described previously (61). Briefly, a HR1 peptide was mixed with a HR2 peptide at a final concentration of 40 μM and incubated at 37°C for 30 min. The mixture was added with Tris–glycine native sample buffer at a ratio of 1 : 1 and then loaded onto a 10 x 1.0-mm Tris-glycine gel (20%) at 25 μl/well. Gel electrophoresis was done with 100V constant voltage at 4 °C for 3 h. The gel was then stained with Coomassie blue and imaged with a Bio-Rad imaging system (Bio-Rad, Hercules, California, USA).

## Acknowledgements

We thank Zene Matsuda at the Institute of Medical Science, University of Tokyo, for providing plasmids and cells for DSP-based cell-cell fusion assay. This work was supported by grants from the National Natural Science Foundation of China (81630061) and the CAMS Innovation Fund for Medical Sciences (2017-I2M-1-014).

## References

1. Wu F, Zhao S, Yu B, Chen YM, Wang W, Song ZG, Hu Y, Tao ZW, Tian JH, Pei YY, Yuan ML, Zhang YL, Dai FH, Liu Y, Wang QM, Zheng JJ, Xu L, Holmes EC, Zhang YZ. 2020. A new coronavirus associated with human respiratory disease in China. Nature 579:265–269.

2. Zhou P, Yang XL, Wang XG, Hu B, Zhang L, Zhang W, Si HR, Zhu Y, Li B, Huang CL, Chen HD, Chen J, Luo Y, Guo H, Jiang RD, Liu MQ, Chen Y, Shen XR, Wang X, Zheng XS, Zhao K, Chen QJ, Deng F, Liu LL, Yan B, Zhan FX, Wang YY, Xiao GF, Shi ZL. 2020. A pneumonia outbreak associated with a new coronavirus of probable bat origin. Nature 579:270–273.

3. Zhu N, Zhang D, Wang W, Li X, Yang B, Song J, Zhao X, Huang B, Shi W, Lu R, Niu P, Zhan F, Ma X, Wang D, Xu W, Wu G, Gao GF, Tan W, China Novel Coronavirus I, Research T. 2020. A Novel Coronavirus from Patients with Pneumonia in China, 2019. N Engl J Med 382:727–733.

4. Perlman S, Netland J. 2009. Coronaviruses post-SARS: update on replication and pathogenesis. Nat Rev Microbiol 7:439–50.

5. Li F. 2016. Structure, Function, and Evolution of Coronavirus Spike Proteins. Annu Rev Virol 3:237–261.

6. Wrapp D, Wang N, Corbett KS, Goldsmith JA, Hsieh CL, Abiona O, Graham BS, McLellan JS. 2020. Cryo-EM structure of the 2019-nCoV spike in the prefusion conformation. Science 367:1260–1263.

7. Walls AC, Park YJ, Tortorici MA, Wall A, McGuire AT, Veesler D. 2020. Structure, Function, and Antigenicity of the SARS-CoV-2 Spike Glycoprotein. Cell doi: 10.1016/j.cell.2020.02.058.

8. Wan Y, Shang J, Graham R, Baric RS, Li F. 2020. Receptor Recognition by the Novel Coronavirus from Wuhan: an Analysis Based on Decade-Long Structural Studies of SARS Coronavirus. J Virol 94.

9. Walls AC, Tortorici MA, Snijder J, Xiong X, Bosch BJ, Rey FA, Veesler D. 2017. Tectonic conformational changes of a coronavirus spike glycoprotein promote membrane fusion. Proc Natl Acad Sci U S A 114:11157–11162.

10. Liu S, Xiao G, Chen Y, He Y, Niu J, Escalante CR, Xiong H, Farmar J, Debnath AK, Tien P, Jiang S. 2004. Interaction between heptad repeat 1 and 2 regions in spike protein of SARS-associated coronavirus: implications for virus fusogenic mechanism and identification of fusion inhibitors. Lancet 363:938–47.

11. Bosch BJ, van der Zee R, de Haan CA, Rottier PJ. 2003. The coronavirus spike protein is a class I virus fusion protein: structural and functional characterization of the fusion core complex. J Virol 77:8801–11.

12. He Y. 2013. Synthesized peptide inhibitors of HIV-1 gp41-dependent membrane fusion. Curr Pharm Des 19:1800–9.

13. Lu L, Liu Q, Zhu Y, Chan KH, Qin L, Li Y, Wang Q, Chan JF, D. L, Yu F, Ma C, Ye S, Yuen KY, Zhang R, Jiang S. 2014. Structure-based discovery of Middle East respiratory syndrome coronavirus fusion inhibitor. Nat Commun 5:3067.

14. Wang C, Xia S, Zhang P, Zhang T, Wang W, Tian Y, Meng G, Jiang S, Liu K. 2018. Discovery of Hydrocarbon-Stapled Short alpha-Helical Peptides as Promising Middle East Respiratory Syndrome Coronavirus (MERS-CoV) Fusion Inhibitors. J Med Chem 61:2018–2026.

15. Bosch BJ, Martina BE, Van Der Zee R, Lepault J, Haijema BJ, Versluis C, Heck AJ, De Groot R, Osterhaus AD, Rottier PJ. 2004. Severe acute respiratory syndrome coronavirus (SARS-CoV) infection inhibition using spike protein heptad repeat-derived peptides. Proc Natl Acad Sci U S A 101:8455–60.

16. Ujike M, Nishikawa H, Otaka A, Yamamoto N, Yamamoto N, Matsuoka M, Kodama E, Fujii N, Taguchi F. 2008. Heptad repeat-derived peptides block protease-mediated direct entry from the cell surface of severe acute respiratory syndrome coronavirus but not entry via the endosomal pathway. J Virol 82:588–92.

17. Liu IJ, Kao CL, Hsieh SC, Wey MT, Kan LS, Wang WK. 2009. Identification of a minimal peptide derived from heptad repeat (HR) 2 of spike protein of SARS-CoV and combination of HR1-derived peptides as fusion inhibitors. Antiviral Res 81:82–7.

18. Aydin H, Al-Khooly D, Lee JE. 2014. Influence of hydrophobic and electrostatic residues on SARS-coronavirus S2 protein stability: insights into mechanisms of general viral fusion and inhibitor design. Protein Sci 23:603–17.

19. Xia S, Yan L, Xu W, Agrawal AS, Algaissi A, Tseng CK, Wang Q, Du L, Tan W, Wilson IA, Jiang S, Yang B, Lu L. 2019. A pan-coronavirus fusion inhibitor targeting the HR1 domain of human coronavirus spike. Sci Adv 5:eaav4580.

20. Zhu Y, Chong H, Yu D, Guo Y, Zhou Y, He Y. 2019. Design and Characterization of Cholesterylated Peptide HIV-1/2 Fusion Inhibitors with Extremely Potent and Long-Lasting Antiviral Activity. J Virol doi: 10.1128/JVI.02312-18.

21. Chong H, Xue J, Zhu Y, Cong Z, Chen T, Wei Q, Qin C, He Y. 2019. Monotherapy with a low-dose lipopeptide HIV fusion inhibitor maintains long-term viral suppression in rhesus macaques. PLoS Pathog 15:e1007552.

22. Zhu Y, Zhang X, Ding X, Chong H, Cui S, He J, Wang X, He Y. 2018. Exceptional potency and structural basis of a T1249-derived lipopeptide fusion inhibitor against HIV-1, HIV-2, and simian immunodeficiency virus. J Biol Chem 293:5323–34.

23. Chong H, Zhu Y, Yu D, He Y. 2018. Structural and Functional Characterization of Membrane Fusion Inhibitors with Extremely Potent Activity against HIV-1, HIV-2, and Simian Immunodeficiency Virus. J Virol 92:e01088–18.

24. Chong H, Xue J, Zhu Y, Cong Z, Chen T, Guo Y, Wei Q, Zhou Y, Qin C, He Y. 2018. Design of Novel HIV-1/2 Fusion Inhibitors with High Therapeutic Efficacy in Rhesus Monkey Models. J Virol 92:e00775–18.

25. Chong H, Xue J, Xiong S, Cong Z, Ding X, Zhu Y, Liu Z, Chen T, Feng Y, He L, Guo Y, Wei Q, Zhou Y, Qin C, He Y. 2017. A Lipopeptide HIV-1/2 Fusion Inhibitor with Highly Potent In Vitro, Ex Vivo, and In Vivo Antiviral Activity. J Virol 91:e00288–17.

26. He Y, Zhou Y, Siddiqui P, Niu J, Jiang S. 2005. Identification of immunodominant epitopes on the membrane protein of the severe acute respiratory syndrome-associated coronavirus. J Clin Microbiol 43:3718–26.

27. He Y, Zhou Y, Wu H, Luo B, Chen J, Li W, Jiang S. 2004. Identification of immunodominant sites on the spike protein of severe acute respiratory syndrome (SARS) coronavirus: implication for developing SARS diagnostics and vaccines. J Immunol 173:4050–7.

28. He Y, Zhou Y, Wu H, Kou Z, Liu S, Jiang S. 2004. Mapping of antigenic sites on the nucleocapsid protein of the severe acute respiratory syndrome coronavirus. J Clin Microbiol 42:5309–14.

29. Cao Z, Liu L, Du L, Zhang C, Jiang S, Li T, He Y. 2010. Potent and persistent antibody responses against the receptor-binding domain of SARS-CoV spike protein in recovered patients. Virol J 7:299.

30. He Y, Barker SJ, MacDonald AJ, Yu Y, Cao L, Li J, Parhar R, Heck S, Hartmann S, Golenbock DT, Jiang S, Libri NA, Semper AE, Rosenberg WM, Lustigman S. 2009. Recombinant Ov-ASP-1, a Th1-biased protein adjuvant derived from the helminth Onchocerca volvulus, can directly bind and activate antigen-presenting cells. J Immunol 182:4005–16.

31. He Y, Li J, Li W, Lustigman S, Farzan M, Jiang S. 2006. Cross-neutralization of human and palm civet severe acute respiratory syndrome coronaviruses by antibodies targeting the receptor-binding domain of spike protein. J Immunol 176:6085–92.

32. He Y, Li J, Jiang S. 2006. A single amino acid substitution (R441A) in the receptor-binding domain of SARS coronavirus spike protein disrupts the antigenic structure and binding activity. Biochem Biophys Res Commun 344:106–13.

33. He Y, Li J, Heck S, Lustigman S, Jiang S. 2006. Antigenic and immunogenic characterization of recombinant baculovirus-expressed severe acute respiratory syndrome coronavirus spike protein: implication for vaccine design. J Virol 80:5757–67.

34. He Y, Li J, Du L, Yan X, Hu G, Zhou Y, Jiang S. 2006. Identification and characterization of novel neutralizing epitopes in the receptor-binding domain of SARS-CoV spike protein: revealing the critical antigenic determinants in inactivated SARS-CoV vaccine. Vaccine 24:5498–508.

35. He Y. 2006. Immunogenicity of SARS-CoV: the receptor-binding domain of S protein is a major target of neutralizing antibodies. Adv Exp Med Biol 581:539–42.

36. He Y, Zhu Q, Liu S, Zhou Y, Yang B, Li J, Jiang S. 2005. Identification of a critical neutralization determinant of severe acute respiratory syndrome (SARS)-associated coronavirus: importance for designing SARS vaccines. Virology 334:74–82.

37. He Y, Lu H, Siddiqui P, Zhou Y, Jiang S. 2005. Receptor-binding domain of severe acute respiratory syndrome coronavirus spike protein contains multiple conformation-dependent epitopes that induce highly potent neutralizing antibodies. J Immunol 174:4908–15.

38. He Y, Jiang S. 2005. Vaccine design for severe acute respiratory syndrome coronavirus. Viral Immunol 18:327–32.

39. He Y, Zhou Y, Siddiqui P, Jiang S. 2004. Inactivated SARS-CoV vaccine elicits high titers of spike protein-specific antibodies that block receptor binding and virus entry. Biochem Biophys Res Commun 325:445–52.

40. He Y, Zhou Y, Liu S, Kou Z, Li W, Farzan M, Jiang S. 2004. Receptor-binding domain of SARS-CoV spike protein induces highly potent neutralizing antibodies: implication for developing subunit vaccine. Biochem Biophys Res Commun 324:773–81.

41. Li W, Moore MJ, Vasilieva N, Sui J, Wong SK, Berne MA, Somasundaran M, Sullivan JL, Luzuriaga K, Greenough TC, Choe H, Farzan M. 2003. Angiotensin-converting enzyme 2 is a functional receptor for the SARS coronavirus. Nature 426:450–4.

42. Li F, Li W, Farzan M, Harrison SC. 2005. Structure of SARS coronavirus spike receptor-binding domain complexed with receptor. Science 309:1864–8.

43. Glowacka I, Bertram S, Muller MA, Allen P, Soilleux E, Pfefferle S, Steffen I, Tsegaye TS, He Y, Gnirss K, Niemeyer D, Schneider H, Drosten C, Pohlmann S. 2011. Evidence that TMPRSS2 activates the severe acute respiratory syndrome coronavirus spike protein for membrane fusion and reduces viral control by the humoral immune response. J Virol 85:4122–34.

44. Shulla A, Heald-Sargent T, Subramanya G, Zhao J, Perlman S, Gallagher T. 2011. A transmembrane serine protease is linked to the severe acute respiratory syndrome coronavirus receptor and activates virus entry. J Virol 85:873–82.

45. Simmons G, Gosalia DN, Rennekamp AJ, Reeves JD, Diamond SL, Bates P. 2005. Inhibitors of cathepsin L prevent severe acute respiratory syndrome coronavirus entry. Proc Natl Acad Sci U S A 102:11876–81.

46. Iwata-Yoshikawa N, Okamura T, Shimizu Y, Hasegawa H, Takeda M, Nagata N. 2019. TMPRSS2 Contributes to Virus Spread and Immunopathology in the Airways of Murine Models after Coronavirus Infection. J Virol 93.

47. Kawase M, Shirato K, van der Hoek L, Taguchi F, Matsuyama S. 2012. Simultaneous treatment of human bronchial epithelial cells with serine and cysteine protease inhibitors prevents severe acute respiratory syndrome coronavirus entry. J Virol 86:6537–45.

48. Zhou Y, Vedantham P, Lu K, Agudelo J, Carrion R, Jr., Nunneley JW, Barnard D, Pohlmann S, McKerrow JH, Renslo AR, Simmons G. 2015. Protease inhibitors targeting coronavirus and filovirus entry. Antiviral Res 116:76–84.

49. Follis KE, York J, Nunberg JH. 2006. Furin cleavage of the SARS coronavirus spike glycoprotein enhances cell-cell fusion but does not affect virion entry. Virology 350:358–69.

50. Kleine-Weber H, Elzayat MT, Hoffmann M, Pohlmann S. 2018. Functional analysis of potential cleavage sites in the MERS-coronavirus spike protein. Sci Rep 8:16597.

51. Park JE, Li K, Barlan A, Fehr AR, Perlman S, McCray PB, Jr., Gallagher T. 2016. Proteolytic processing of Middle East respiratory syndrome coronavirus spikes expands virus tropism. Proc Natl Acad Sci U S A 113:12262–12267.

52. Hoffmann M, Kleine-Weber H, Schroeder S, Kruger N, Herrler T, Erichsen S, Schiergens TS, Herrler G, Wu NH, Nitsche A, Muller MA, Drosten C, Pohlmann S. 2020. SARS-CoV-2 Cell Entry Depends on ACE2 and TMPRSS2 and Is Blocked by a Clinically Proven Protease Inhibitor. Cell doi: 10.1016/j.cell.2020.02.052.

53. Shirato K, Kanou K, Kawase M, Matsuyama S. 2017. Clinical Isolates of Human Coronavirus 229E Bypass the Endosome for Cell Entry. J Virol 91.

54. Shirato K, Kawase M, Matsuyama S. 2018. Wild-type human coronaviruses prefer cell-surface TMPRSS2 to endosomal cathepsins for cell entry. Virology 517:9–15.

55. Baron SA, Devaux C, Colson P, Raoult D, Rolain JM. 2020. Teicoplanin: an alternative drug for the treatment of coronavirus COVID-19? Int J Antimicrob Agents doi: 10.1016/j.ijantimicag.2020.105944:105944.

56. Zhang J MX, Yu F, Liu J, Zou F, Pan T, Zhang H. Teicoplanin potently blocks the cell entry of 2019-nCoV. BioRxiv 2020:20200205935387 https://doi.org/10.1101/2020.02.05.935387.

57. Wang M, Cao R, Zhang L, Yang X, Liu J, Xu M, Shi Z, Hu Z, Zhong W, Xiao G. 2020. Remdesivir and chloroquine effectively inhibit the recently emerged novel coronavirus (2019-nCoV) in vitro. Cell Res 30:269–271.

58. Martinez MA. 2020. Compounds with therapeutic potential against novel respiratory 2019 coronavirus. Antimicrob Agents Chemother doi: 10.1128/AAC.00399-20.

59. Gautret P, Lagier JC, Parola P, Hoang VT, Meddeb L, Mailhe M, Doudier B, Courjon J, Giordanengo V, Vieira VE, Dupont HT, Honore S, Colson P, Chabriere E, La Scola B, Rolain JM, Brouqui P, Raoult D. 2020. Hydroxychloroquine and azithromycin as a treatment of COVID-19: results of an open-label non-randomized clinical trial. Int J Antimicrob Agents doi: 10.1016/j.ijantimicag.2020.105949:105949.

60. Zhu Y, Ding X, Yu D, Chong H, He Y. 2019. The Tryptophan-Rich Motif of HIV-1 Gp41 Can Interact with the N-Terminal Deep Pocket Site: New Insights into the Structure and Function of Gp41 and Its Inhibitors. J Virol doi: 10.1128/JVI.01358-19.

61. Zhang X, Ding X, Zhu Y, Chong H, Cui S, He J, Wang X, He Y. 2019. Structural and functional characterization of HIV-1 cell fusion inhibitor T20. AIDS 33:1–11.

